# Does a map-level insulation constraint improve sequence-based prediction of 3D genome folding? A Stage 1 Registered Report with a reverse-complement-equivariant state-space model (HiFold-Mamba)

**DOI:** 10.64898/2026.10.24.761502

**Authors:** Mohammad Malekpouri, Somayeh Lotfi

## Abstract

**Background:** Sequence-to-contact-map models are trained with pixel-wise losses, although the features of interest (insulation, domain boundaries, loops) are summaries of the map. Insulation-based losses help Hi-C resolution enhancement. No pre-specified test has asked whether they improve *sequence-based* prediction, judged against annotations independent of the training data.

**Proposed study:** HiFold-Mamba predicts 448 × 448 maps at 2,048 bp from 1,048,576 bp of DNA on the DNALongBench contact-map task. Every component is reverse-complement-equivariant: convolutional tower, bidirectional Mamba blocks, multi-scale mixer and symmetric pairwise head. We pre-register three hypotheses:

- **H1:** it exceeds *each* of three trained baselines (reproduced Akita, a parameter-matched Transformer and a parameter-matched CNN) by ≥ 0.01 stratum-adjusted correlation (SCC).
- **H2:** an insulation and boundary loss, computed from the predicted map with the operator used on the observed map, improves boundary AUROC and loop AP against independent 4DN boundary calls and ENCODE CTCF ChIA-PET loops, at equal training steps and without loss of SCC.
- **H3:** pretraining plus few-shot adaptation beats training from scratch on four held-out cell lines.

Tests use a paired bootstrap over 237 one-megabase test blocks (one-sided α = 0.025).

**Pilot data:**

- **Verification**. The benchmark target, the window coordinates and the network’s equivariance were verified.
- **Locus-shared reference**. Copying other cell lines’ measured maps at the same locus matches published Akita on the benchmark’s SCC (0.227 vs. 0.230).
- **SCC weightings**. Two weightings in common use weight distance strata in opposite directions and agree only moderately (ρ = 0.54).
- **Comparison rule**. In simulations calibrated on the pilot data, a per-block-best comparison against several baselines has no power. We therefore test against each baseline, which gives ≥ 90 % power at a true gain of 0.02 when between-model correlation is ≥ 0.95.
- **H2 positive control**. On validation windows, observed maps recover the independent boundaries (AUROC 0.61–0.91) and loops (0.86–0.90). This fixed the insulation window and showed that these annotations are largely shared across cell types.

## 1. Introduction

### 1.1 Background

Chromosomes fold into compartments, topologically associating domains (TADs) and CTCF-anchored loops, and disruption of this folding by structural variants can rewire gene regulation (Suhas S. P. Rao et al. 2014; Krietenstein et al. 2020). Hi-C and Micro-C measure folding genome-wide but are expensive per cell type and condition. Models that predict contact maps directly from DNA sequence therefore serve two purposes: they test how much of folding is encoded in sequence, and they allow in-silico perturbation of that sequence.

Sequence-only models now span several scales. Akita predicts 448 × 448 maps at 2,048 bp from 1,048,576 bp of sequence with a convolutional tower and a 2D head (Fudenberg, Kelley, and Pollard 2020). DeepC uses megabase-scale transfer learning (Schwessinger et al. 2020). Orca predicts from kilobase to whole-chromosome scale (Zhou 2022). AlphaGenome predicts contact maps among thousands of tracks from 1 Mb of sequence (Avsec et al. 2026). Models that add cell-type-specific inputs, for example C.Origami (sequence with CTCF and accessibility) (Tan et al. 2023) and Hi-Compass (sequence and a generalized CTCF profile, with chromatin accessibility as the only cell-type-specific input) (Sun et al. 2026), address cross-cell-type prediction, which sequence alone cannot.

### 1.2 What is not known

#### (i) Whether objectives defined on map-level structure help sequence-based prediction

The sequence-to-map models whose objectives we could verify, Akita and the DNALongBench baselines, are trained with a pixel-wise mean squared error (Fudenberg, Kelley, and Pollard 2020; Cheng et al. 2025). The biologically important outputs are summaries of the map: insulation profiles, boundary positions and loop calls. Structure-aware losses exist in neighbouring problems. VEHiCLE, a Hi-C resolution-enhancement network, adds an insulation-vector loss and a loss on its derivative (a “delta” vector marking boundaries) (Highsmith and Cheng 2021). Hi-Compass adds a structural-similarity (SSIM) term to an accessibility-based predictor (Sun et al. 2026). Two things are untested: whether such terms help when the input is sequence alone and the target is a distance-normalized log observed/expected (O/E) map, and whether any gain survives evaluation against boundaries and loops that were not derived from the training data. A model whose insulation term is scored with boundaries called from its own training maps is graded partly on what it was told to fit.

#### (ii) How strong current sequence-only predictions are, relative to what a locus alone explains

On DNALongBench’s contact-map task, Akita reaches SCC 0.230 averaged over its five cell lines, and HyenaDNA and Caduceus 0.115–0.133 (Cheng et al. 2025) (Table 4 there). Contact maps are partly shared across cell types. How much of the published accuracy is cell-type-specific is not reported. It matters for interpreting any gain (Section 3.3).

#### (iii) Whether published differences are statistically resolved

Benchmark results are typically means over windows, without a test that respects the overlap and spatial correlation of genomic windows. A recent study imputing 4DN contact maps found its model beat the strongest simple baseline in only 8 of 14 test cases (Min et al. 2025). That is a reminder that simple references are hard to beat and that gains must be tested, not asserted.

### 1.3 This study

We register a design that answers (i), with (ii) and (iii) built into the analysis. The test vehicle is HiFold-Mamba, an RC-equivariant bidirectional state-space model. We do **not** claim the encoder family as a contribution: bidirectional, RC-equivariant Mamba layers for DNA are the core of Caduceus (Schiff et al. 2024), and a bidirectional Mamba has been applied to chromatin-interaction prediction from sequence and accessibility (ChiMamba) (Zhang et al. 2024); we position ChiMamba from its published abstract, as the full chapter was not accessible to us. At 512 bins per window, attention is computationally cheap, so we make **no efficiency claim**. The encoder matters to this study only because the matched Transformer and CNN ablations (H1, Section 2.4) separate encoder choice from the structural objective (H2).

Contributions, conditional on the pre-registered outcomes:

1. A **pre-registered test (H2) of a map-level insulation and boundary constraint** for sequence-based contact-map prediction. The constraint is computed from the predicted map with exactly the operator applied to the observed map, adapted from the insulation-loss idea of VEHiCLE (Highsmith and Cheng 2021) to a log-O/E target. It is evaluated on **independent** boundary and loop annotations, with the accessions fixed in this protocol.
2. An **RC-equivariant network end to end**: tower, state-space blocks, multi-scale mixer, pair-wise head, FiLM cell adapter and per-cell distance bias. f(RC(x), c) = rot_180_ f(x, c) holds by construction and is verified numerically to 10^−4^ in fp32 (pilot, Section 3.2).
3. A **statistical design** for this benchmark: a paired block bootstrap with an intersection–union comparison against every trained baseline, a simulation showing that a per-block-maximum comparison (our original plan) has no power at this sample size, locus-shared references kept outside the comparison set, and conditional claims fixed in advance (Section 2.15).
4. Pilot measurements of independent interest: a non-learned locus-shared reference that matches published Akita on the benchmark metric, and a moderate disagreement between two SCC weightings in common use (Section 3.3).

## 2. Methods

### 2.1 Data

#### Benchmark

DNALongBench contact-map task (Cheng et al. 2025), Harvard Dataverse doi:10.7910/DVN/AZM25S (CC0), downloaded 2026-10-21 to 2026-10-23, every file MD5-verified against the Dataverse record. Genome build hg38. Inputs are 1,048,576 bp windows (one-hot). Targets are 448 × 448 maps at 2,048 bp after cropping 32 bins per side, stored as the upper triangle with diagonal offset 2 (99,681 values per map and cell line).

#### Training cell lines (five)

Training cell lines (five).
| Line | Assay (source) | Akita/DNALongBench split<br>(train / valid / test windows) |
| --- | --- | --- |
| HFF | Micro-C (Krietenstein et al. 2020); 4DN 4DNFIP5EUOFX | 7,008 / 419 / 413 |
| H1-hESC | Micro-C (Krietenstein et al. 2020); 4DN 4DNFI1O6IL1Q | same windows |
| GM12878 | in situ Hi-C, MboI (Suhas S. P. Rao et al. 2014) | same windows |
| IMR-90 | in situ Hi-C, MboI (Suhas S. P. Rao et al. 2014) | same windows |
| HCT116 | in situ Hi-C, unsynchronized (Suhas S. P. Rao et al. 2017) | same windows |

#### Held-out cell lines (H3 only; four)

HAP1, HeLa, HepG2 and K562, which DNALongBench added from 4DN with its own windows (7,009 / 6,342 / 6,291) on the same 354 virtual contigs. We checked that none of their validation or test windows overlaps an Akita training window, and none of Akita’s validation or test windows overlaps one of their training windows (Section 3.1). These lines are never used in training or pretraining.

#### Split

The official files: random non-overlapping virtual contigs, assigned 8:1:1 (Fudenberg, Kelley, and Pollard 2020; Cheng et al. 2025). No window-level random split is used anywhere. A chromosome-held-out split is a separate robustness analysis (Section 2.13).

#### Window coordinates

The TFRecords carry no coordinates. Record *i* corresponds to window *i* of Akita’s sequences.bed. We confirmed this by matching the stored one-hot sequence to hg38 for sampled records in every split (agreement > 0.99 in every sampled record; Section 3.1). Coordinates are needed for the genomic blocks of the bootstrap and for the structural annotations.

### 2.2 Target space

The target is the benchmark’s processed map, used unchanged. From Akita’s processing code (`akita_data_read.py`), the five training lines are ICE-balanced coolers at 2,048 bp, adaptively coarse-grained, divided by local expected (O/E), natural-log transformed, clipped to [−2, 2], with missing bins interpolated, the near-diagonal band set to 0 and a 2D Gaussian (σ = 1) applied. We confirmed in the files that values lie in [−2, 2], that no NaN or mask is shipped, and that no training, validation or test map is all-zero (Section 3.1). The four held-out lines were processed with `kernel_stddev 0`, i.e. **without** smoothing (`their options.json`). We use them unchanged and state this as a limitation. It does not bias Δ_H3, because both H3 arms see identical targets, but absolute SCC on these lines is not comparable with the training lines.

No log or other transform is applied to the target or to predictions in the contact loss. Any transform inside a structural operator is applied identically to prediction and target (Section 2.6).

### 2.3 Model

Let x ∈ {A,C,G,T}^L with L = 1,048,576, and c a cell-line index. Channels-last features h ∈ ℝ^(L×2D) carry the RC action R(h) = flip_length(flip_channel(h)). With one-hot input in A, C, G, T order, R is exactly the reverse complement. Every layer is RC-equivariant (f ∘ R = R ∘ f), using Caduceus-style parameter sharing (Schiff et al. 2024): RCPS(F)(h) = [F(h), R(F(R h))].

- **Conv tower**. RC-equivariant stem convolution (kernel 15, 64 channels), then 11 blocks of RC convolution (kernel 5), RC LayerNorm (tied affine parameters [g, flip(g)]), GELU and maxpool 2. Channels 64–256. This takes 2^20 bp to 512 bins, followed by an RC-equivariant projection to d = 256. The tower is shared by all cell lines and runs once per window.
- **State-space blocks**. 8 pre-norm residual blocks. Each applies RCPS around a projection to 128 channels followed by a bidirectional Mamba-1 layer (Gu and Dao 2023) (forward and reversed scans, summed; d_state 16, expansion 2), then an RC-equivariant feed-forward sublayer (×2). The selective scan is a chunked, compiled pure-PyTorch implementation, tested against a sequential reference.
- **FiLM cell adapter**. A learned embedding e_c ∈ ℝ^32 is linearly mapped to (γ, β), applied as h ← h + (1 + tie(γ)) ⊙ Mixer(LN(h)) + tie(β), where tie(v) = [v, flip(v)] keeps the modulation RC-symmetric (Perez et al. 2018). Its initialization is zero.
- **Multi-scale mixer**. The states after blocks 2, 5 and 8 are each average-pooled at 1, 2, 4 and 8 bins, upsampled, projected with RC-equivariant linear maps, summed and normalized.
- **Pairwise head**. With Z the mixed representation, J the channel flip, and P, Q ∈ ℝ^(d×64): U = [ZP, (JZ)P] and V = [ZQ, (JZ)Q], giving

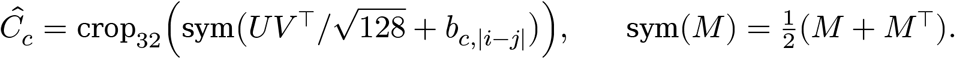 The pairing gives J M J = M for the bilinear term, so RC of the input rotates the map by 180°. The distance bias b_{c,|i−j|} is learned **per cell line**. The pilot targets show line-specific residual distance profiles after O/E normalization (trough −0.17 to −0.30), which a shared bias cannot express. A new held-out line gets its own bias, learned during fine-tuning.
- **Size**. 3,322,560 parameters (paper configuration). The development-only 8 GB fallback configuration is never reported.

#### Equivariance precondition

Unit tests assert f(R x, c) = rot_180_ f(x, c) in fp32 (absolute tolerance 10^−4^, relative 10^−5^) for every component and for the full network. The full network is tested at full input length, on random and real sequences, in evaluation and deterministic training mode, with and without gradient checkpointing. A looser bf16 check is also run, and a negative control (an unpaired head) must fail. All pass (Section 3.2).

### 2.4 Comparison set *ℬ* and references

*ℬ*, the set of trained sequence models against which H1 is tested, is fixed here:

| Model in $\mathcal{B}$ | Definition | Parameters |
| --- | --- | --- |
| Akita (reproduced) | PyTorch port of the published architecture with Keras semantics (pre-activation residual blocks, BatchNorm momentum and $\varepsilon$ as in Keras, dilation rates 1, 2, 3, 5, 9, 16, 29, 50, mean outer product + distance channel, symmetrized 2D dilated residual stack), trained with the published optimizer (SGD, lr 0.0065, momentum 0.99575, clip 10, batch 2) and RC + $\pm 11$ bp shift augmentation | 746,149 |
| Matched Transformer | HiFold with each Bi-Mamba replaced by multi-head self-attention over bins with a learned per-head bias on $\vert i - j\vert $, in the same RCPS wrapper (RC-equivariant) | 3,340,992 (+0.55 %) |
| Matched CNN (“strong CNN”) | HiFold with each Bi-Mamba replaced by a residual dilated 1D CNN (6 layers, dilation $2^k$, width 73), in the same RCPS wrapper | 3,319,720 (−0.09 %) |

The two matched models share the conv tower, FiLM adapter, mixer, head, data, optimizer, schedule and checkpoint rule with HiFold-Mamba, so they isolate the sequence mixer.

#### Not in *ℬ*

- **Caduceus-Ph/PS and HyenaDNA**. Their published DNALongBench scores (SCC 0.115–0.133 (Cheng et al. 2025)) are far below Akita’s, which is in *ℬ*. Fine-tuning them at 1 Mb context is outside our compute budget (Section 4). They appear in the results only as published context.
- **Non-learned references, reported for context and never in** *ℬ*. (a) The *global cell-line mean*: the mean training map of the line, the same prediction for every window. (b) The *locus-shared mean*: the mean of the **other** four training lines’ measured maps at the same window. (c) The same with a per-line offset. Because (b) and (c) use measured test-region maps, they are oracle-style references for how cell-type-shared a map is, not sequence predictors.

Every retrained model receives the same EMA (decay 0.999), checkpoint-selection rule and tuning budget (no tuning beyond the published or matched settings). The Akita reproduction is checked against its published per-line scores (Cheng et al. 2025) on the benchmark metric. It counts as reproduced if its five-line mean test SCC (benchmark implementation) is within 0.02 of the published 0.230. A larger gap is reported as a deviation, with the published values shown alongside, and the reproduced Akita stays in *ℬ* either way.

### 2.5 Training

AdamW (weight decay 0.01), learning rate 3 × 10^−4^, 1,000-step warmup, cosine decay. bf16 autocast, with the selective scan in fp32. Gradient checkpointing; batch 1 window × 5 lines with 8-step accumulation (effective 8 windows). 70,080 steps, i.e. 10 passes over 7,008 windows. RC-equivariant models need no RC augmentation. All models use ±11 bp shift augmentation with N padding. **Checkpoint selection:** EMA weights with the highest mean validation SCC (HiCRep weighting, Section 2.7), evaluated every 7,008 steps. “HiFold-Mamba” in H1, H3 and the ablations means the *Base* arm (contact loss only, Section 2.6), selected this way. For H2 only, both arms are compared at their **final** EMA weights (step 70,080). Otherwise validation selection could return a Struct checkpoint from before the structural terms were switched on, which would make H2 null by construction. Three seeds (0, 1, 2) per model. Every run logs its configuration, configuration hash and git commit.

### 2.6 Structural objective (H2)

With 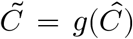 and 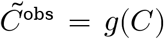, where *g*(*x*) = exp(2tanh(*x*/2)) (smooth, about exp near 0, bounded to [e^−2^, e^2^], the clip range of the target), the crossing-square insulation of bin *i* is

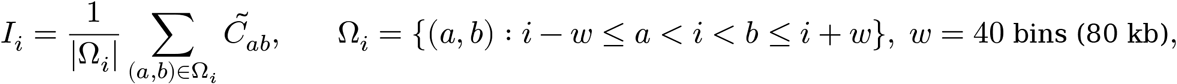

defined for i ∈ [w, 448 − w). The window w = 40 was fixed on validation data, where it recovered the independent boundaries better than the originally planned w = 20 in every line (Section 3.4). It is computed with an integral image, which makes it exactly differentiable (Crane et al. 2015). The boundary strength is the depth of the log-insulation dip relative to its neighbourhood, 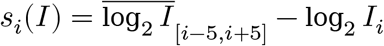. Then

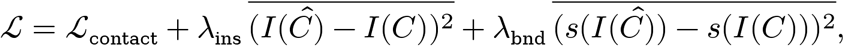

with *ℒ*_contact_ the masked MSE over the 99,681 upper-triangle values. Because the operator is linear in and g is smooth, the structural gradient reaches the map itself. The loss can only be reduced by changing the predicted map, unlike an auxiliary structure head, whose output can be fitted independently of the map. The constraint is partial: different maps can share an insulation profile.

#### Schedule (equal optimizer steps and data)

The *Base* arm trains with ℒ_contact_ for all 70,080 steps. The *Struct* arm is identical, with the same seeds, up to step 52,560, and then adds the structural terms for the remaining 17,520 steps. Total optimizer steps and data are identical. The structural terms add two integral images and a few elementwise operations per 448 × 448 map, which is small next to the network’s forward and backward pass. Their cost is measured and reported.

#### Choice of λ (pre-registered rule)

On a 1-seed pilot (seed 0) scored on **validation windows only**, we fork the Base run at step 52,560 and train the Struct phase for each (λ_ins, λ_bnd) ∈ {0.1, 0.3, 1.0}^2^. Among settings whose validation SCC is within 0.005 of the Base pilot, we select the one with the highest validation AUROC against the independent boundaries (Section 2.8). The selection uses the same boundary datasets as the H2 evaluation, but only in validation regions, which are disjoint from the test regions. Ties go to the smaller λ_ins + λ_bnd. If no setting meets the SCC condition, we take the smallest pair. The selected pair is then fixed for all seeds and for the test evaluation.

#### No loop term in v1

An earlier internal plan included a loop loss. We drop it, because its only usable positives would come from the same ChIA-PET datasets used to evaluate loops, which would make H2’s loop criterion partly self-supervised. H2’s loop criterion therefore tests whether insulation and boundary constraints *transfer* to loops.

### 2.7 Metrics

**Primary: SCC with HiCRep weights** (Yang et al. 2017), per window and cell line. For strata d = 2, …, 447 (diagonal offset 2, as in the benchmark), we compute the Pearson correlation r_d between predicted and true values on the d-th diagonal (N_d values). We combine them as SCC = Σ_d w_d r_d / Σ_d w_d, with 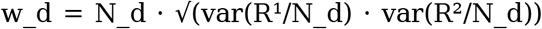, where R^1^ and R^2^ are the within-stratum ranks (HiCRep’s variance-stabilized weights; ties averaged). There is no 2D smoothing (h = 0), because the target is already smoothed. Strata with fewer than two values or a constant side are skipped. A window with no scorable stratum has an undefined SCC (Section 2.9, exclusions).

#### Secondary (always reported next to the primary)

SCC with the benchmark’s released implementation (`scripts/metrics.py::scc`) for comparability with published numbers. It sets w_d = N_d · var(0, …, N_d − 1)/N_d^4^ ≈ 1/(12 N_d), so weight *decreases* with stratum size and long-distance strata dominate. HiCRep weights grow ≈ N_d/12, so short-distance strata dominate. Also reported: Pearson correlation (PCC) per window; SCC per distance band (10–100 kb, 100–500 kb, 0.5–0.92 Mb); MSE. Spearman correlation, if reported, is labelled as such.

### 2.8 Structural evaluation data (independent of the training targets)

Accessions were fixed and data downloaded (MD5-verified) before any model was trained:

| Line | Boundaries (4DN processed boundaries, GRCh38) | Loops (ENCODE CTCF ChIA-PET loops, GRCh38) |
| --- | --- | --- |
| HFF | 4DNFI3CGVRKU, in situ Hi-C, FA+DSG, DpnII (Akgol Oksuz et al. 2021), 5 kb. Same lab as the Micro-C target; different assay and study | none available |
| H1-hESC | 4DNFICEHILOK, Zhong lab, untreated control (Calandrelli et al. 2023), 5 kb | ENCFF401IWZ (ENCSR095SXC, Lee lab, JAX) |
| GM12878 | 4DNFIQBWXNL1, Ren lab dilution Hi-C (Selvaraj et al. 2013), 10 kb | ENCFF780PGS (ENCSR184YZV, Ruan lab, JAX) |
| IMR-90 | 4DNFIUI2RM2Q, Ren lab dilution Hi-C (Jin et al. 2013), 10 kb | ENCFF682YFU (ENCSR076TTY, Ruan lab, JAX) |
| HCT116 | 4DNFIG49IPR6, Cremins lab, untreated WAPL-mAID2 cells (Emerson et al. 2022), 5 kb | ENCFF431BUN (ENCSR278IZK, Lee lab, JAX; unmodified HCT116) |

None of these datasets is the source of a training target. For four of the five lines they come from a different laboratory.

- **Boundaries**. We use the shipped 4DN boundary calls as the primary set, with no caller of our own. Their calling method (an insulation-score pipeline) is of the same family as our operator, which we state as a limitation. A bin is a positive if its 2,048 bp interval overlaps a called boundary interval (Strong and Weak calls alike) extended by one bin on each side. All other scorable bins are negatives. The score is s_i of the **predicted** map (Section 2.6). A window scores bins i ∈ [w + 5, 448 − w − 5). Test windows overlap, so each genomic bin is one item: its score is the mean of s_i over every test window that scores it, then over seeds.
- **Loops**. Positives are ChIA-PET PET clusters with **PET count ≥ 10** whose anchors’ mid-points fall in two different output bins inside the same window, at separation 20 kb–915 kb. Negatives are all other bin pairs within the same windows, matched to positives on separation in 10 log-spaced distance bins (five negatives per positive). The score is the predicted map value at the pair, averaged over every test window that contains both bins, then over seeds. Each genomic pair is one item.
- **Secondary, assay-orthogonal set (reported, not tested)**. ENCODE CTCF ChIP-seq peaks containing a CTCF motif.
- **Internal consistency only**. Boundaries called from the observed DNALongBench maps by our own operator.

### 2.9 Statistical analysis

All confirmatory tests use one procedure:

1. Score each test window, cell line, model and seed. Average over the five lines, then over the three seeds.
2. Assign each window to the 1 Mb genomic block containing its midpoint (237 blocks in the test set) and average within blocks. Blocks, not windows, are the resampling unit, because windows overlap and neighbouring loci are correlated.
3. For H1: Δ_{m,b} = S_ours,b − S_m,b separately for **each** m ∈ *ℬ* (intersection–union test, below).
4. Paired bootstrap over blocks, 10,000 resamples, using the same resamples for every comparison. One-sided test of H_0_: E[Δ] ≤ 0 at α = 0.025, with the 95% percentile interval. Its lower limit is the 97.5% one-sided bound. The p-value is (1 + #{Δ* ≤ 0}) / (B + 1).

#### Criteria

Criteria.
| Hypothesis | Success requires |
| --- | --- |
| H1 | against **every** $m \in \mathcal{B}$: mean $\Delta_m \geq +0.01$ **and**<br>lower bound $> 0$. An intersection-union test,<br>so the level is $\alpha$ with no correction<br>(stats.against_each). It replaces a<br>per-block-maximum rule that the power<br>analysis showed to be unpassable (Section<br>2.10) |
| H2 | **all three hold** (intersection-union, so no multiplicity correction): (a) Struct – Base independent-boundary AUROC $\geq +0.02$ with lower bound $> 0$. AUROC is computed per line over that line’s items (Section 2.8), then averaged over the five lines. Each bootstrap resample draws blocks with replacement, shared across lines and arms. An item enters with weight equal to the number of times its block (that of its bin, or of the pair’s midpoint) was drawn, and AUROC is recomputed exactly under these weights<br>(`structure_eval.paired_block_bootstrap_metric`).<br>(b) Loop AP $\geq +0.02$, same procedure, averaged over the four lines with loop data; (c) SCC non-inferiority: lower bound of $\Delta$ SCC(Struct – Base) $> -0.005$ |
| H3 | $\Delta_{H3} = \text{SCC}(\text{pretrain} + \text{few-shot}) - \text{SCC}(\text{scratch})$, per held-out line and limited regime (10 %, 25 %, 50 %), Holm-corrected over the three regimes within a line. Success requires a positive one-sided bound in at least one regime in **at least two of the four** lines |
| Multi-scale (N-M) | H1 met **and** $\Delta$ SCC(full – no-mixer ablation) has a lower bound $> 0$ |

A per-window Wilcoxon signed-rank test is reported as a secondary, anti-conservative check only. **Outcome-neutral checks (positive controls), fixed in advance**.

- **Pipeline**. The equivariance tests pass at the frozen commit.
- **Akita reproduction**. Our reproduced Akita reaches the criterion of Section 2.4. If it does not, we report this and keep it in *ℬ*, so a weak reproduction cannot make H1 easier, because H1 must beat every baseline.
- **Sanity floor**. Every trained model must beat the global cell-line mean (one-sided block bootstrap). A model that fails is reported as a failed run and diagnosed (data, target, split) before any hypothesis test that uses it.

#### Exclusions

Windows and lines with an undefined SCC (no scorable stratum) are excluded from that model’s mean and listed. None exist in the pilot references. **Missing maps:** a target map that is exactly zero everywhere is Akita’s placeholder for a region absent from the source cooler. It is treated as missing (NaN) in every loss and metric. Only 27 of 7,009 K562 training maps are affected (held-out lines; none in any validation or test split, and none for the five training lines). Bins without a defined boundary score, and loop pairs outside every test window, are not items. No other exclusion is permitted.

#### Block-size sensitivity

every confirmatory test is repeated with 3 Mb blocks. Windows straddle 1 Mb block borders, so residual dependence may make 1 Mb intervals slightly narrow. Disagreements are reported but do not change the decision.

#### The margins and their scale

The H1 margin of 0.01 is on the primary (HiCRep) scale, where the locus-shared reference scores 0.53. It is the smallest gain we would consider worth reporting: about 2 % of the reference level, and similar to the spread between published foundation-model variants on this task (Cheng et al. 2025). The H2 margins (+0.02 AUROC and AP) are set at the same order of magnitude relative to their chance levels.

#### Error control across hypotheses

H1, H2, H3 and N-M support distinct claims, each with its own conclusion and title rule (Section 2.15). Each is tested at one-sided α = 0.025 with no correction across hypotheses. Within hypotheses, H1 and H2 are intersection–union tests (level α by construction), H3 is Holm-corrected within each line, and N-M is gated by H1 (a fixed-sequence test). Every outcome is reported whatever its direction, so no claim depends on choosing among tests after the fact.

### 2.10 Sample size and power

The test set is fixed by the benchmark (413 windows in 237 blocks), so the question is the sensitivity of the design, not the sample size. We simulated block means for our model and the baselines with the between-block standard deviation observed in the pilot (0.193, HiCRep SCC of the locus-shared reference averaged over lines). Models were equicorrelated with ρ ∈ {0.8, 0.9, 0.95, 0.98}, and we applied the full H1 rule (2,000 simulations × 2,000 bootstrap resamples; `scripts/power_analysis.py`). Single-comparator power:

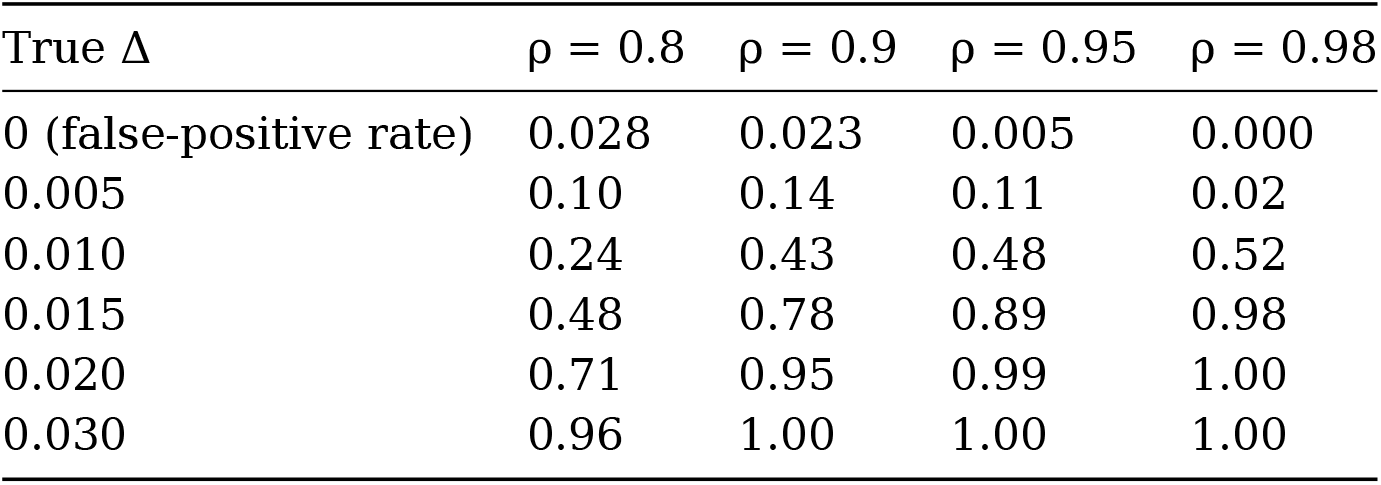

The false-positive rate stays at or below α. Because H1 requires the **point estimate** to reach +0.01, power at a true effect of exactly 0.01 cannot exceed about 0.5.

#### Three baselines

We first simulated the rule originally planned, Δ_b = S_ours,b − max_m S_m,b, with three baselines that are each worse than ours by the true Δ. Its power was **0.00 for every Δ ≤ 0.02 and every ρ**, and 0.08 at Δ = 0.03 (ρ = 0.98) (results/pilot/power_h1_max3.tsv). Most blocks hold one or two windows. The maximum of three noisy block means therefore exceeds their common mean by about 0.85 σ(1 − ρ)^½, i.e. 0.02–0.07 SCC across the simulated ρ, which is larger than any realistic gain. We replaced it with the intersection–union rule. Its worst-case power, with three equally strong baselines, is:

| True $\Delta$ (vs. each) | $\rho = 0.8$ | $\rho = 0.9$ | $\rho = 0.95$ | $\rho = 0.98$ |
| --- | --- | --- | --- | --- |
| 0 | 0.001 | 0.003 | 0.001 | 0.000 |
| 0.010 | 0.08 | 0.19 | 0.26 | 0.24 |
| 0.015 | 0.22 | 0.56 | 0.76 | 0.94 |
| 0.020 | 0.49 | 0.88 | 0.99 | 1.00 |
| 0.030 | 0.91 | 1.00 | 1.00 | 1.00 |

When the baselines differ in strength, only the strongest constrains the result, and power approaches the single-comparator table. The design therefore has ≥ 0.9 power for a true advantage of about 0.02 over every baseline when ρ ≥ 0.95 (Fig. 2c). The paired-model correlation ρ is unknown before training. It will be estimated on **validation** windows from the Akita reproduction and the first HiFold seed, and reported with the power table, before any test evaluation. We keep the 0.01 margin, set as the smallest gain worth reporting, and state openly that a non-significant result at that size is inconclusive, not evidence of no effect.

**Figure 1:**
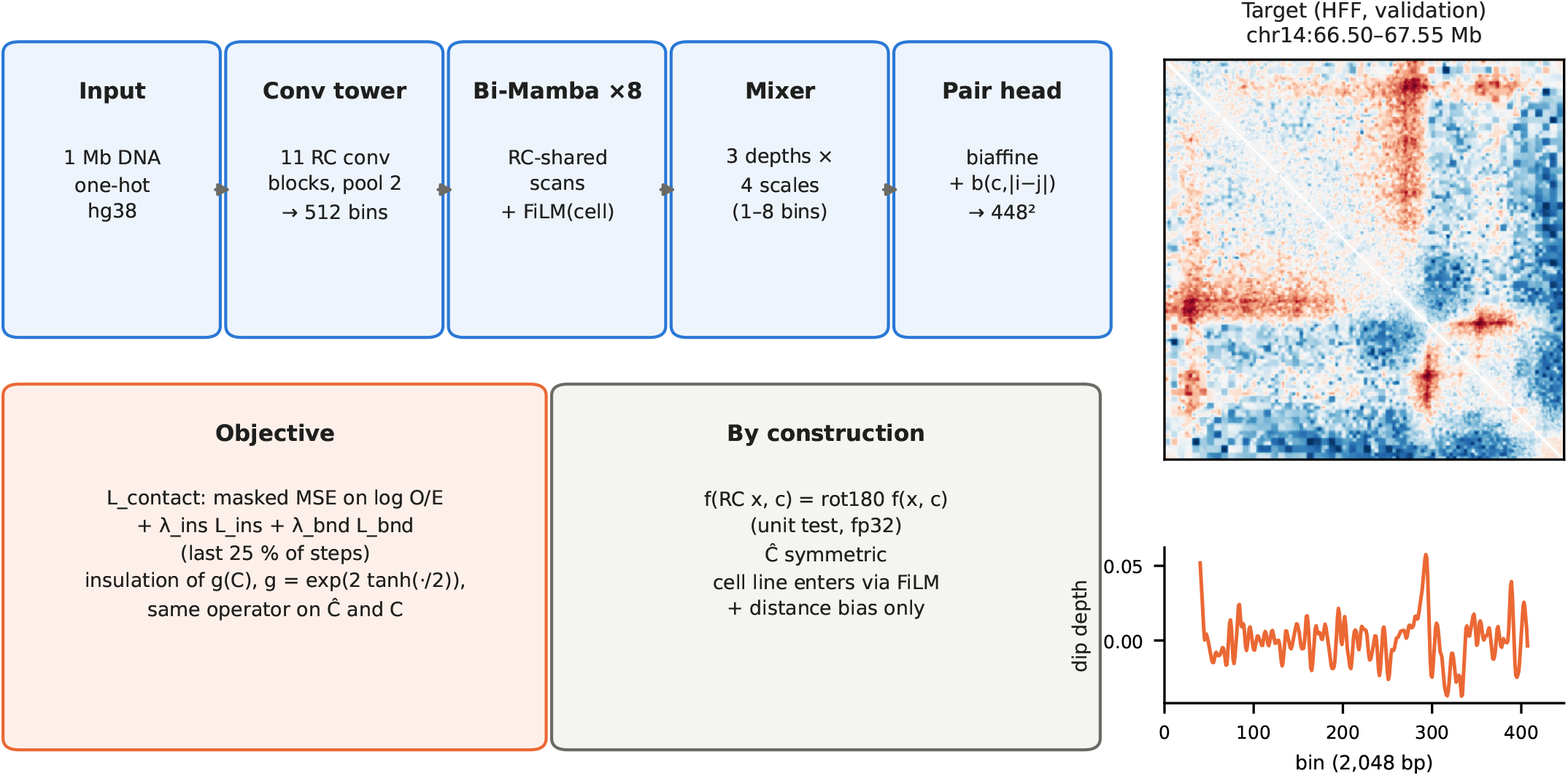
HiFold-Mamba and the structural objective. Left: architecture (every component RC-equivariant) and the training objective. Right: a validation target map (HFF) and the boundary-strength profile s_i computed from it with the operator used in the loss and in evaluation.

**Figure 2:**
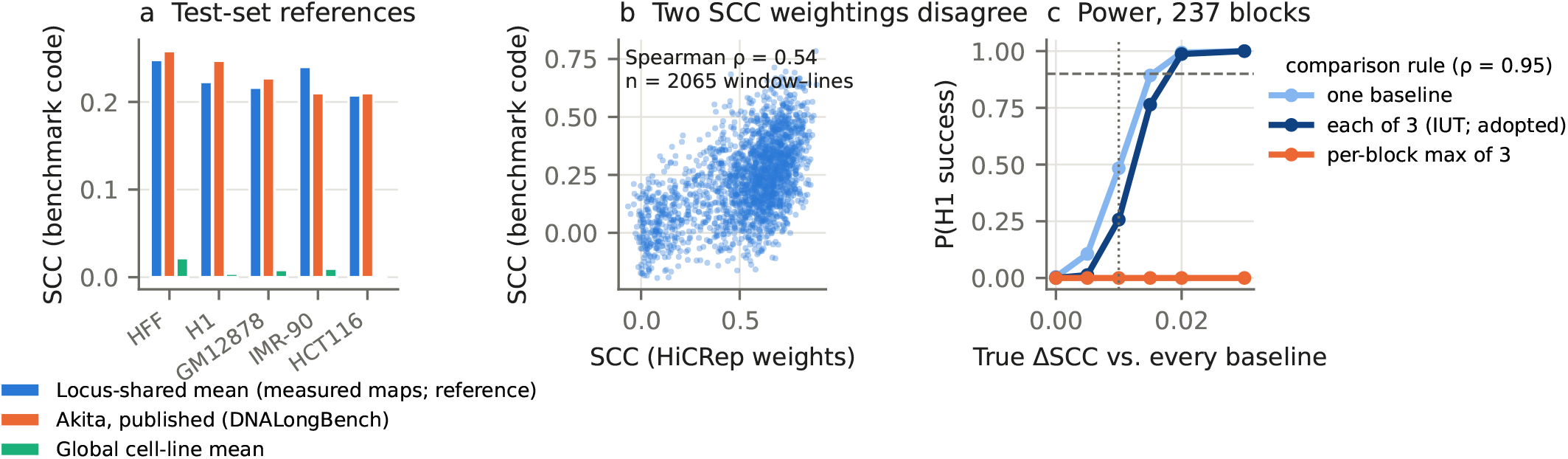
Pilot data on the test set (no trained model). **a**, Per-line SCC (benchmark implementation) of the locus-shared reference (measured maps of the other four lines) against published Akita (Cheng et al. 2025) and the global cell-line mean. **b**, Per-window agreement of the two SCC weightings for the locus-shared reference. **c**, Simulated probability that the H1 rule succeeds as a function of the true ΔSCC (ρ = 0.95; 237 blocks). Three comparison rules are shown: a single baseline; each of three equally strong baselines (intersection–union, adopted); and the per-block maximum of three, which never succeeds. Dashed line: 0.9; dotted line: the 0.01 margin.

### 2.11 H3: few-shot adaptation to held-out lines

For each held-out line (HAP1, HeLa, HepG2, K562), the **pretrain arm** starts from the selected five-line HiFold-Mamba Base checkpoint (seed-matched) with a new cell embedding and a new distance-bias row, and fine-tunes all weights. The **scratch arm** has the same architecture with random initialization. Both arms use identical data and identical optimizer steps: 3,504 steps, batch 1, 8-step accumulation, learning rate 1 × 10^−4^ with 200-step warmup and cosine decay. Fine-tuning data are contiguous-block subsets of the line’s official training windows: the 354 virtual contigs are ordered by a fixed random permutation (seed 0), and whole contigs are added until 10 %, 25 % or 50 % of the training windows are covered, so the subsets are nested and identical across arms and seeds. The full training set is also used (reported, not counted toward success). Evaluation uses HiCRep SCC on the line’s **official test windows**, aggregated into 1 Mb blocks (these windows overlap at 32 kb stride), with the Section 2.9 bootstrap over that line’s blocks. Three seeds per arm. Read-depth-limited regimes are out of scope (they would need reprocessing from read pairs). **Zero-shot reference (secondary, not a criterion)**. The pretrained model with the held-out line’s embedding set to the mean of the five training embeddings and no fine-tuning. Reporting Δ(pretrain + few-shot − zero-shot) shows whether adaptation adds anything beyond the pretrained model’s prior. Equal-compute comparison with scratch is the conventional test of transfer, but it can favour pretraining by design.

### 2.12 H4: structural variants

H4 is confirmatory only if at least ten structural variants with published before/after contact maps in a compatible assay are assembled **before** Stage 2 analysis. The criterion is the sign agreement of predicted vs. observed change in insulation at the variant, tested with a one-sided binomial test at α = 0.025. Otherwise H4 is reported as an in-silico analysis (deletions and inversions of CTCF sites and boundaries), labelled exploratory, with no claim about measured biological effects.

### 2.13 Robustness analysis (secondary)

A chromosome-held-out split (test chr1, chr8, chr12; validation chr3, chr6; windows re-derived from Akita’s coordinates) is trained with one seed for HiFold-Mamba and each model in *ℬ*. It is reported separately, never pooled with the primary results and never used for any confirmatory decision.

### 2.14 Ablations (each 3 seeds, same schedule; reported with the §2.9 procedure, exploratory except N-M)

- Bi-Mamba → matched Transformer and → matched CNN (these are the *ℬ* members).
- –**RC equivariance**. Same layer shapes and FLOPs, with each reverse half given its own parameters and RC augmentation used instead. This has 6.64 M parameters, twice the full model’s, so it favours the ablation.
- –**multi-scale mixer**. The pair head sees only the final state at full resolution (2.96 M parameters).
- Struct vs. Base (H2), and each structural term alone (λ_bnd = 0 or λ_ins = 0).
- Insulation window w ∈ {20, 80} (one seed; Struct arm only).

### 2.15 Interpretation, conditional claims and stopping rules

The final title and claims follow from the outcomes, fixed now:

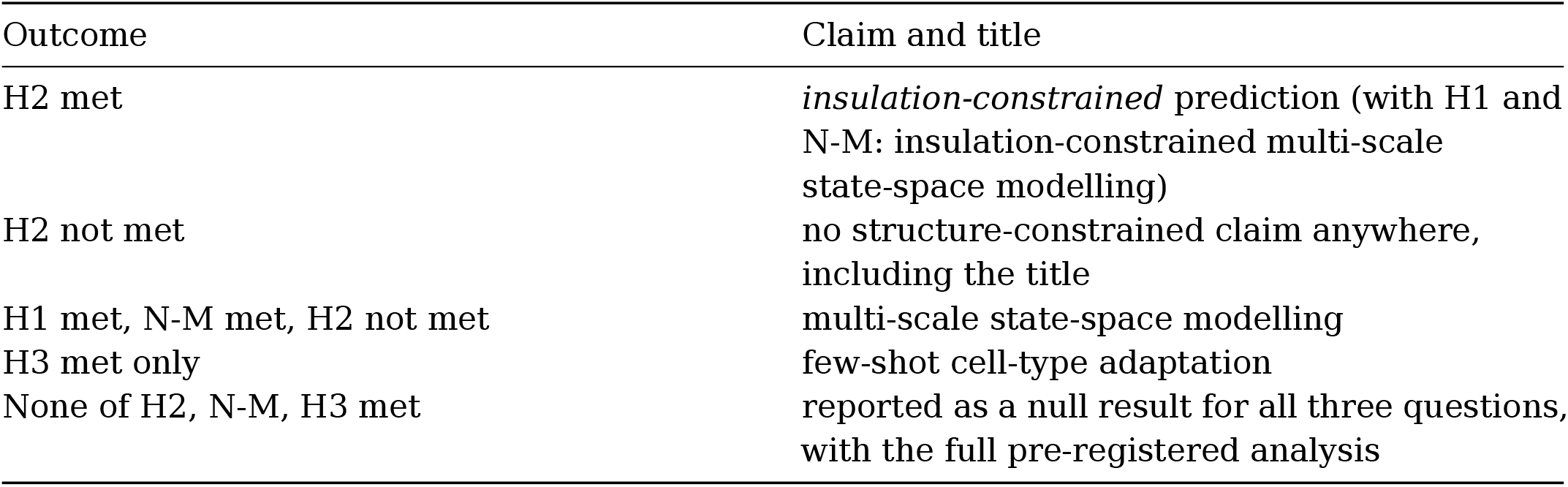

We never claim efficiency at 512 bins, “multi-megabase” scale, physical or energy-based modelling, or RC-equivariant bidirectional Mamba as a contribution in itself. The locus-shared references are never a stopping condition. The global cell-line mean is a sanity floor only. The pilot shows it scores SCC ≈ 0 in this target space (Section 3.3), so it cannot discriminate between models.

### 2.16 Data access and bias control

Data observed before this protocol was frozen:

- Training and validation targets (used for development, profiling, the λ and ρ pilots).
- Test-set **targets**, used only for non-learned references (global mean, locus-shared mean), the SCC-variant comparison and the power analysis. **No trained model has been evaluated on the test set**.
- The independent boundary and loop files. They were intersected only with **observed validation maps** for the positive control (Section 3.4), never with a model prediction or a test window.

Safeguards:

- Test-set scoring of trained models is disabled until Stage 1 acceptance. All model selection uses validation data.
- The evaluation code is frozen at the commit recorded at submission. Any later change is listed with its reason in the Stage 2 report.
- The Akita reproduction is trained before acceptance, on training and validation windows only. Its validation scores are pilot data.

#### Bias-control level (PCI RR taxonomy)

We declare **Level 2**. The benchmark data exist, and some of them, including the test-set targets, have been accessed and partly observed by the authors (above). The key variables that answer H1–H3 have not been observed and do not yet exist: the trained models’ test-set predictions, and their intersection with the independent annotations on test windows. They will be generated only after in-principle acceptance. The safeguards above are the additional steps to strengthen bias control that this level requires.

### 2.17 Study design table

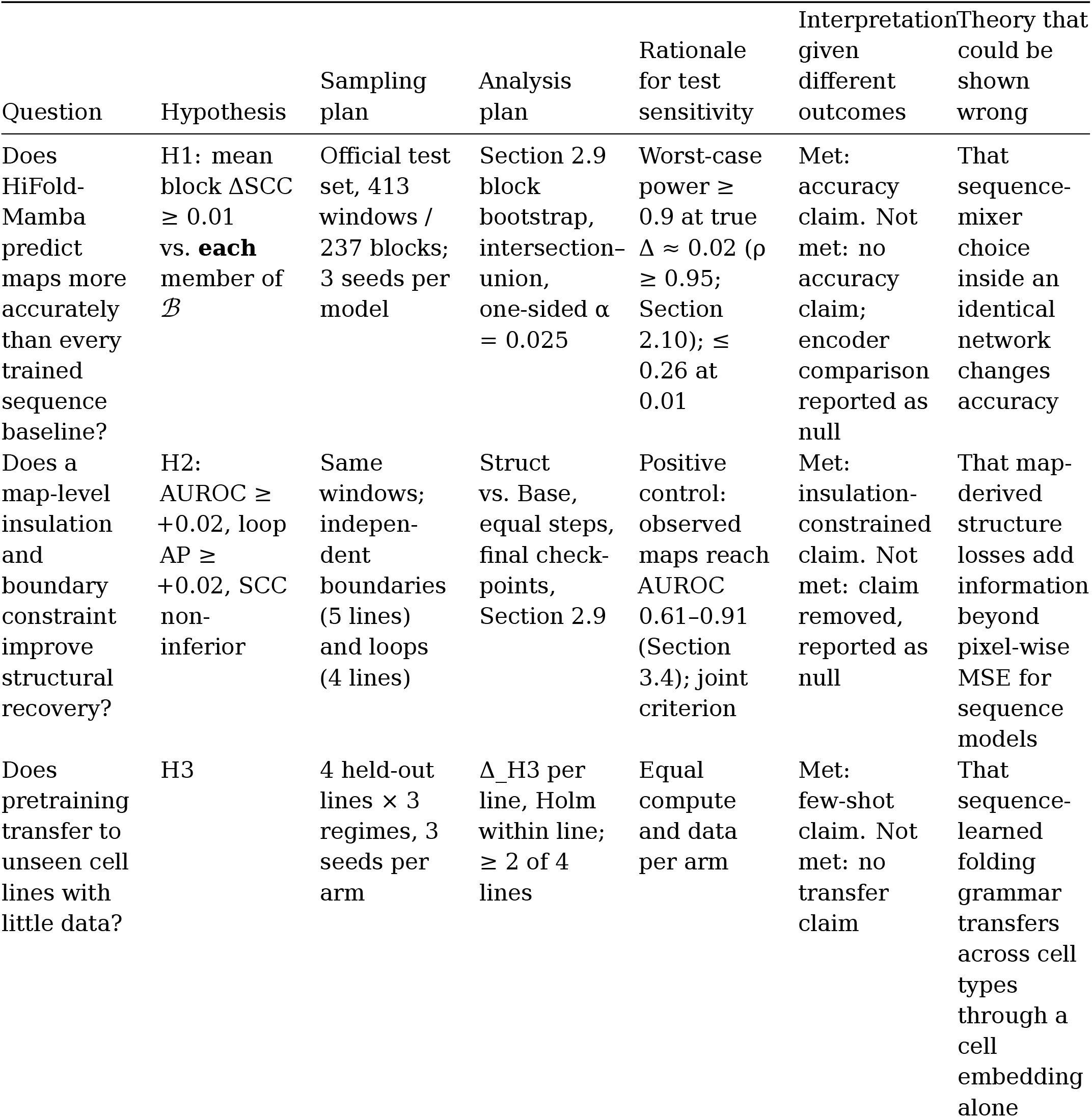
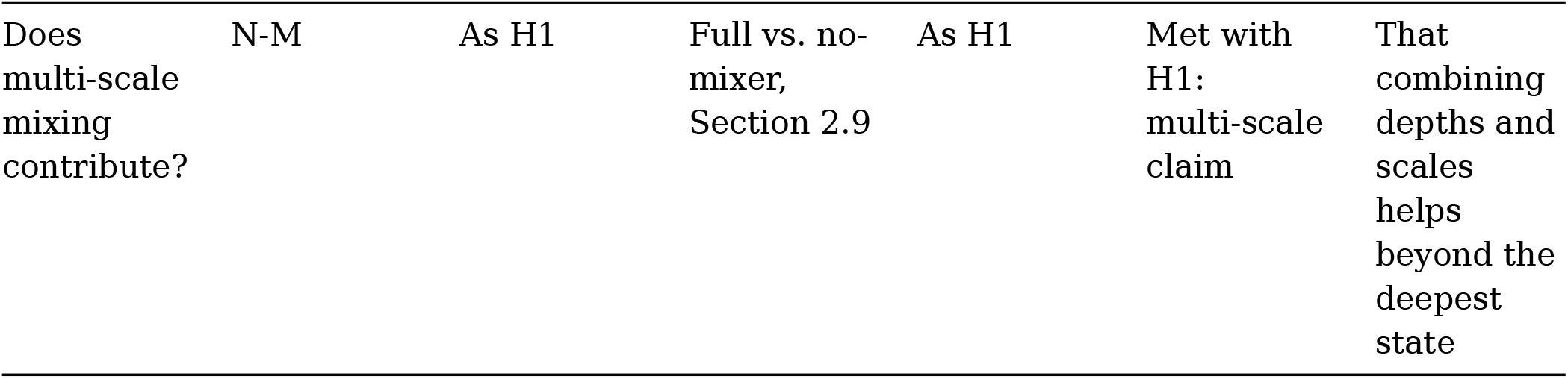

## 3. Pilot data

All pilot numbers come from logged scripts in the repository. None involves a trained model on the test set.

### 3.1 Data verification

- **Target transform (G1)**. Confirmed against Akita’s processing code and the files, as described in Section 2.2. Shipped values lie in [−2, 2]; no NaN; zero all-zero maps among 7,008 / 419 / 413 windows in any of the five lines.
- **Coordinates**. For sampled records in every split (74 train, 6 validation, plus the test set), the stored one-hot sequence matches hg38 at the window’s coordinates with > 99 % identity. Record order equals window order.
- **Held-out lines**. Cached 7,009 / 6,342 / 6,291 windows, with sampled records matching hg38 in every split. 27 all-zero (missing) K562 training maps are handled as described in Section 2.9.
- **Split hygiene**. Zero overlap between the held-out lines’ validation and test windows and Akita’s training windows, and zero overlap in the reverse direction (scripts/check_h3_overlap.py).
- **Residual distance profiles**. Each line has its own residual P(s) after O/E normalization, with troughs of −0.17 to −0.30. This motivated the per-cell distance bias, which a synthetic-data training test confirms is needed to fit line-specific profiles (tests/test_train_smoke.py).

### 3.2 Implementation checks

- **Equivariance**. All equivariance tests pass in fp32 (10^−4^ / 10^−5^), including the full-length network on real sequence. The negative control (unpaired head) is detected. The − RC ablation is verified to break equivariance while keeping the outputs symmetric. In total, 64 unit tests pass at the protocol commit, including the H1 intersection–union test and the exact weighted AUROC/AP used for H2.
- **Metrics**. The vectorized benchmark SCC reproduces the released loop implementation exactly. HiCRep SCC is tested against a direct implementation.
- **Compute** (RTX 5050, 8 GB, bf16, one window × five lines): HiFold-Mamba 1.34 s per step at 3.84 GiB peak; matched Transformer 0.50 s; matched CNN 0.47 s; Akita 0.25 s per step at batch 2 (4.73 GiB).
- **Akita reproduction (training and validation only)**. Seed 0 completed the full 70,080-step schedule. Final validation SCC (mean over the five lines, EMA weights): 0.196 with the benchmark code, 0.469 with HiCRep weights, PCC 0.396. Learning was slow until about step 45,000 and was still rising at the final step. The run passes our internal pre-use check (validation SCC, benchmark code, ≥ 0.15); the reproduction criterion of Section 2.4 is assessed on the test set at Stage 2. Validation and test values are not directly comparable. Median step time 0.27 s when the GPU was not shared; wall-clock time 14.8 h because the GPU was shared with another project.

### 3.3 Reference baselines on the test set

| Reference (not in $\mathcal{B}$) | SCC, HiCRep weights | SCC, benchmark code | PCC |
| --- | --- | --- | --- |
| Global cell-line mean | 0.014 | 0.009 | 0.089 |
| Locus-shared mean<br>(other four lines’<br>measured maps) | 0.531 | 0.227 | 0.582 |
| *Akita, published*<br>*(Cheng et al. 2025)* | — | 0.230 | — |

The published value is compared under the assumption that DNALongBench computed it with the scc function of its released code, which we use verbatim. The paper reports SCC without giving the implementation.

Three findings shape the design (Fig. 2):

1. **The global mean is uninformative in this target space**. After O/E normalization, a line’s average map is nearly flat within every distance stratum, so its SCC is ≈ 0. “Beating the global mean” is therefore only a sanity floor, not a meaningful bar.
2. **A non-learned, locus-only reference matches published Akita on the benchmark metric**. Per line: HFF 0.248 vs. 0.258, H1 0.223 vs. 0.247, GM12878 0.216 vs. 0.227, IMR-90
3. 0.240 vs. 0.210, HCT116 0.207 vs. 0.210 (Fig. 2a). Much of the measurable signal is shared across cell types at a locus. The reference uses measured maps, so it is not a competitor. But it shows that benchmark-level accuracy alone does not establish cell-type-specific prediction, which is why H3 and the per-line analyses are part of the design.
4. **The two SCC weightings disagree moderately** (Spearman ρ = 0.54 across 2,065 window– line pairs; Fig. 2b). The two weightings point in opposite directions: ∝ 1/N_d (benchmark; long-range strata dominate) vs. ∝ N_d (HiCRep; short-range strata dominate). The two scores therefore emphasize different parts of the map, and model rankings can depend on the variant. We pre-register HiCRep as primary and report both.

### 3.4 Positive control for the H2 evaluation (validation windows, no model)

Before any model is trained, the H2 pipeline must show that **observed** maps recover the independent annotations. Otherwise a null H2 could not be told apart from a broken evaluation. We ran the exact Stage 2 code (item deduplication, labelling, AUROC and AP) on the 419 validation windows. For each line we scored its own observed map and the locus-shared reference (`scripts/pilot_structure.py`; Fig. 3).

**Figure 3:**
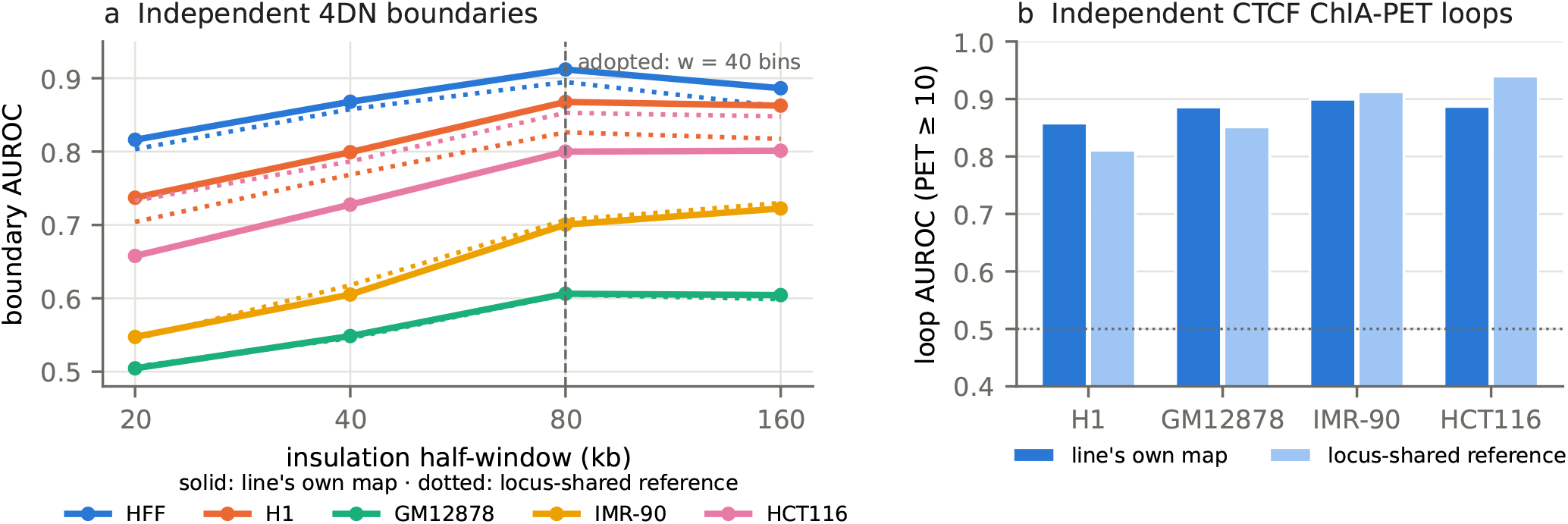
Positive control for the H2 evaluation on validation windows (no model). **a**, AUROC of the boundary score s_i against independent 4DN boundaries, as a function of the insulation window, for each line’s own observed map (solid) and the locus-shared reference (dotted). **b**, AUROC of the observed map value at independent CTCF ChIA-PET loops (PET ≥ 10) against distance-matched negatives.

- **Boundaries** (≈ 110,000 genomic bins per line; 3.5–5.6 % positive). With the observed map, AUROC at w = 40 is 0.91 (HFF), 0.87 (H1), 0.80 (HCT116), 0.70 (IMR-90) and 0.61 (GM12878). Every line improves from w = 20 to w = 40 (by 0.04–0.10), and w = 80 adds nothing (Fig. 3a). We therefore fixed w = 40 for the loss and the evaluation, with w = 20 and 80 as sensitivity analyses. The two lines with weak recovery are the ones whose independent calls come from shallow dilution Hi-C at 10 kb. Their positional and calling noise lowers the attainable AUROC, and the line-average in H2 inherits it.
- **Loops** (PET ≥ 10, 5 distance-matched negatives per positive). The observed maps rank ChIA-PET pairs above matched negatives with AUROC 0.86–0.90 and AP 0.71–0.79 (chance 0.17).
- **Annotations are largely shared across cell types**. The locus-shared reference recovers the boundaries almost as well as each line’s own map, and better for HCT116 (0.85 vs. 0.80). It also recovers the loops better for IMR-90 and HCT116. H2 therefore measures recovery of *largely invariant* folding features, not cell-type specificity. We interpret it that way, and cell-type-specific claims rest on H3 alone.
- **Scale of the margins**. The attainable boundary AUROC (0.61–0.91) leaves room for the +0.02 margin in every line. In loops, 0.02 AP is about 3 % of the observed-map level.

## 4. Timeline and compute

On the authors’ RTX 5050 (8 GB), the confirmatory programme needs about:

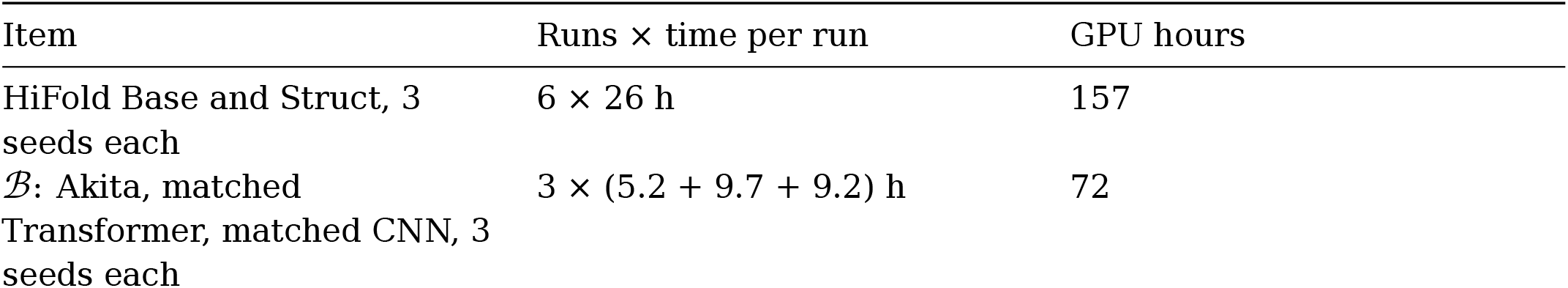
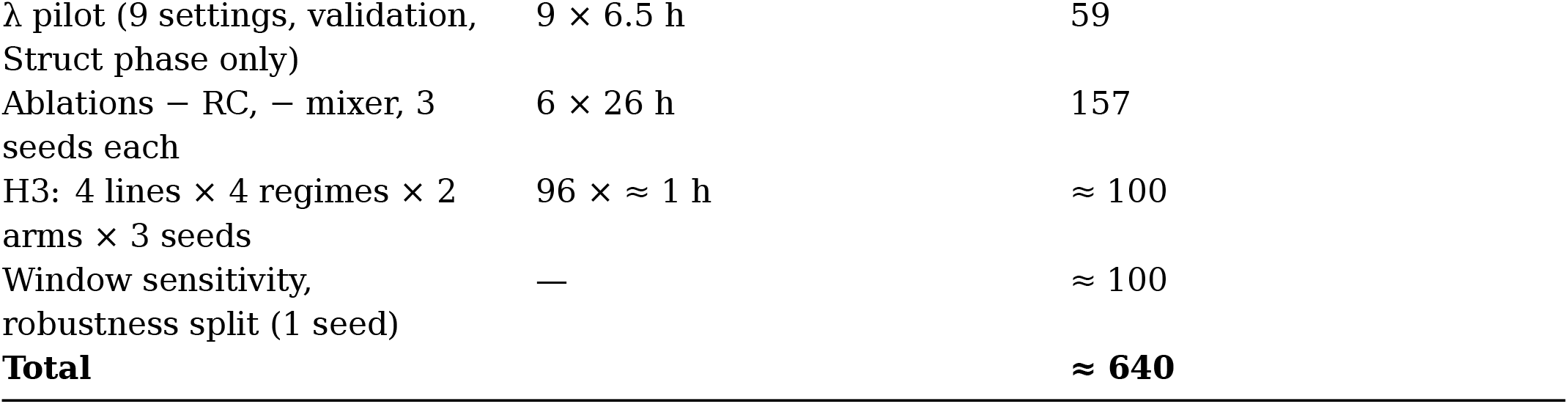

That is about four weeks of continuous single-GPU time. Per-run times are measured step times (Section 3.2) multiplied by the schedule; the Akita time is the measured median for seed 0 without GPU sharing. Stage 2 submission is planned within six months of in-principle acceptance.

## 5. Limitations (stated before outcomes)

1. **Scale**. About 1 Mb windows; megabase-scale only. Orca and AlphaGenome cover longer contexts.
2. **Sequence only**. Without cell-type-specific inputs, prediction for an unseen line needs adaptation data (H3). Zero-shot transfer would require epigenomic inputs.
3. **Annotation independence is partial**. The independent boundaries come from an insulation-family caller, and for HFF from the same laboratory as the training target. Some boundary sets are shallow dilution Hi-C at 10 kb. HFF has no independent loop set.
4. **Held-out lines are unsmoothed**, so their absolute SCC is not comparable with the training lines.
5. **Sensitivity**. H1 is under-powered at exactly the 0.01 margin, more so against three equally strong baselines (Section 2.10). A non-significant H1 at that size is inconclusive.
6. **Benchmark split**. Random virtual contigs. The chromosome-held-out analysis is secondary.
7. **Encoder precedent**. RC-equivariant bidirectional Mamba follows Caduceus; the structural constraint itself adapts VEHiCLE’s insulation loss to a new setting.

## Statements

### Data availability

All data are public: DNALongBench (doi:10.7910/DVN/AZM25S, CC0); 4DN boundary files and ENCODE ChIA-PET files with the accessions in Section 2.8. Checksums and access dates are in `docs/data_manifest.md`.

### Code availability

https://github.com/malekpouri/HiFold-Mamba: model, baselines, training, evaluation, statistics, tests and configurations. Environment: Python 3.12.14, PyTorch 2.14.1 (CUDA 13.0), NumPy 2.5.4, SciPy 1.18.1, pinned in `requirements.lock.txt` and `uv.lock`. The selective scan is a pure-PyTorch implementation, so no custom CUDA kernels are needed. Code licence: MIT.

### Computational resources

All work used one workstation: NVIDIA GeForce RTX 5050 GPU (8 GB, Blackwell), Intel Core i5-11400 CPU (6 cores, 12 threads), 32 GB RAM, Ubuntu Linux; bf16 mixed precision with gradient checkpointing. No cloud or HPC resources were used. Stage 1 compute: data preparation, reference baselines, power simulations and the H2 positive control ran on the CPU; the only GPU training was the Akita reproduction (seed 0, 14.8 h wall-clock on a shared GPU). Stage 2 needs about 640 GPU-hours on the same hardware (Section 4).

### Author contributions (CRediT)

Mohammad Malekpouri: Conceptualization, Methodology, Software, Investigation, Formal analysis, Data curation, Visualization, Project administration, Writing – original draft, Writing – review & editing. Somayeh Lotfi: Writing – review & editing. Both authors approved the final manuscript.

### Use of AI tools

Claude (Anthropic) was used to generate code and to draft and edit the manuscript text. The authors made or approved every design decision and take full responsibility for the content.

